# Genetic structure of mesophotic and shallow *Acropora aculeus* on isolated atolls of Eastern Australia

**DOI:** 10.1101/2025.04.08.647879

**Authors:** A Hernández-Agreda, JA Hoey, D van Hulten, P Hernandez, L Peplow, P Muir, O Hoegh-Guldberg, MJH van Oppen, P Bongaerts

**Affiliations:** California Academy of Sciences, San Francisco, California, USA; School of Biological Sciences, University of Auckland, Auckland, New Zealand; Australian Institute of Marine Science, Townsville, Queensland, Australia; Global Change Institute and School of Biological Sciences, The University of Queensland, Saint Lucia, Australia; School of Biosciences, The University of Melbourne, Parkville, Victoria, Australia

## Abstract

*Acropora* is the most diverse and widespread coral genus in the world. Although known for its critical ecological role in shallow-water habitats, its abundance and diversity at upper mesophotic depths have only recently been uncovered. Consequently, little is known about the genetic structuring of mesophotic *Acropora* populations and their potential ecological and evolutionary relationships with shallow populations. Here, we present the first population genomic evaluation of the depth-generalist coral *Acropora aculeus* to assess genetic structuring across depths (10 and 40 m) and regions (the Great Barrier Reef (GBR) and the Western Coral Sea (WCS)). We observed strong geographic differentiation between regions, indicating the relative isolation of WCS atolls, with some gene flow occurring from WCS to the GBR, but rarely in the opposite direction. Conversely, we observed no geographic or depth-related genetic structuring within regions, although sample sizes limited the evaluation of local allelic patterns of depth structuring. Our results indicate that widespread connectivity within regions characterizes the broadcast spawning *A. aculeus*. The lack of depth differentiation should be explored to see whether the wide depth range of some species may be safeguarded against a rapidly changing climate.

## Introduction

*Acropora* is the most speciose, common, and widespread genus of reef-building corals in the world (Renema et al., 2016; Wallace, 1999). Accounting for roughly 30% of scleractinian coral species (Veron, 2000; Wallace, 1999), the genus includes over 400 nominal species (Hoeksema & Cairns, 2025), although recent genetic work has suggested a systematic revision is required (Bridge et al., 2024; Cowman et al., 2020). Staghorn species of the *Acropora* genus dominate tropical and subtropical shallow reef habitats, providing the three-dimensional structural complexity that supports reef biodiversity (Ball et al., 2021a; Veron, 2000; Wallace, 1999). Furthermore, *Acropora* dominance historically defined the ecological zonation of shallow reef habitats – reef flats, crests, and fore reef zone – across the Caribbean and the Indo-Pacific, and this zonation is known to be driven by factors such as light penetration, hydrodynamic energy, and oxygen content (Muir, Wallace, Done, et al., 2015; Renema et al., 2016; Wallace, 1999). However, due to the significant decline of *Acropora* in the Caribbean (Jackson et al., 2014), this ecological role has largely been lost in that bioregion (Alvarez-Filip et al., 2009; Gladfelter, 1982; Perry et al., 2015). The occurrence and ecological role of *Acropora* at mesophotic depths has rarely been acknowledged despite its relative abundance at upper mesophotic depths (∼30-50 m in the Indo-Pacific (Colin & Lindfield, 2019; Denis et al., 2019; Englebert et al., 2015, 2017; Longenecker et al., 2019; Montgomery et al., 2019; Muir, Wallace, Bridge, et al., 2015; Sinniger et al., 2019)). Although several *Acropora* species are deep-water specialists (e.g., *A. pichoni* and *A. tenella*), ∼10-20% of *Acropora* species in eastern Australia are reported to be common at both shallow and mesophotic depths (Muir, Wallace, Bridge, et al., 2015).

Given the dominant ecological role of the *Acropora* genus, understanding its taxonomic boundaries, connectivity, and genetic variation within and among populations has become critical for the effective management and conservation of coral reefs. However, species delimitation and the study of meta-population structure within the genus have been hampered by introgressive hybridization, morphological plasticity, cryptic diversity, and limitations of traditional sequence markers for species identification (Richards et al., 2013, 2016; Richards & Hobbs, 2015; Van Oppen et al., 2001; Vollmer & Palumbi, 2004; Wallace, 1999). Although *Acropora’s* evolutionary success in the Indo-Pacific is partially attributed to the pervasive introgressive hybridization (Veron, 2000; Willis et al., 2006), its permeable taxonomic boundaries create complex networks of intermittently interbreeding species (known as syngameons (Ladner & Palumbi, 2012; Richards et al., 2013)) and extensive morphological variation (Bridge et al., 2024; Van Oppen et al., 2001). Furthermore, widespread *Acropora* species in the Indo-Pacific exhibit significant cryptic diversity. High levels of genetic differentiation detected between cryptic lineages residing in separate reefs and ocean basins suggest no contemporary gene flow and likely represent distinct nominal species that might have been synonymized (Ball et al., 2021b; Richards et al., 2016). Despite the growing evidence of cryptic diversity and the substantial proportion of depth-generalist species within the genus, no assessments have yet explored the occurrence of genetic depth differentiation in *Acropora*.

Eastern Australia is principally known for the Great Barrier Reef (GBR). Further offshore is the lesser-known Western Coral Sea (WCS), which hosts remote atolls surrounded by deep oceanic water (Bongaerts et al., 2011; Ceccarelli, 2011) and highly diverse reefs (Bridge et al., 2019; Ceccarelli, 2011). While ecologically distinct from the GBR (Hoey et al., 2020), it is hypothesized that WCS reefs play a stepping-stone role in connectivity between the Western Pacific and the GBR (Bode et al., 2006). Additionally, mesophotic coral ecosystems (MCEs, >30 m) are extensive in the WCS, with atoll walls extending to the lower mesophotic zone, where coral has been recorded as deep as 125 m (Englebert et al., 2017; Muir et al., 2018) and reef habitats are dominated by scleractinian corals, octocorals, and the macroalga *Halimeda* (Bongaerts et al., 2011; Galbraith et al., 2024). Recently, atolls of the WCS have suffered major disturbance events, and their remoteness raises questions about the availability of external recruitment sources for recovery.

Here, we assess the genetic structure of the depth-generalist coral *Acropora aculeus* (Dana & Dana, 1846) using reduced representation genome sequencing (nextRAD) on samples collected from both shallow (10 m) and upper mesophotic (40 m) reefs in the Great Barrier Reef (GBR) and the Western Coral Sea (WCS). Our results reveal significant divergence between GBR and WCS bioregions but no differentiation related to depth or locations (within regions). We argue that the potential for panmixis of *A. aculeus* across depth (within bioregions) warrants further investigation into the refuge role of mesophotic populations on isolated reefs like those in the WCS.

## Methods

### Sample collections

Small fragments (1-5 cm) of *Acropora aculeus* (*n* = 181) were collected via SCUBA diving in eight locations of the Great Barrier Reef and Western Coral Sea in 2012 (Fig. 2A). Coral fragments were sampled at shallow (10 m) and upper mesophotic (40 m) depths with a minimum of 3 m of separation between colonies to minimize the potential for collecting clones. Using bone cutters, small tissue samples were subsampled from the main fragment, preserved in a salt-saturated buffer solution (20% DMSO/0.5 M EDTA), and stored at -20 °C until further processing. Following preservation, skeletal fragments were bleached (10% bleach), rinsed, and dried for morphological examination. Specimens were examined using a microscope and identified according to Wallace (1999). Several specimens of the species *A. cerealis* were present in our collection, as well as specimens with non-typical morphologies, which were considered *Acropora* cf. *aculeus* in the analysis. This collection was part of the “XL Catlin Seaview Survey” under permits G12/35281.1, and G14/37294.1 (Great Barrier Reef Marine Park Authority), 018-CZRS1207626-01, 018-RRRW-131031-01 (Australian Department of the Environment).

### Library preparation, sequencing, and clustering

Genomic DNA was extracted using a modified “Wayne’s method” (see Bongaerts et al., 2017). Samples quality and yield were evaluated through gel electrophoresis and Qubit fluorometer (Invitrogen), and normalized before library preparation. Libraries were prepared using the nextRAD method by SNPsaurus LLC. In brief, fragmented gDNA was ligated with adapter sequences using Nextera reagents (Illumina Inc) and PCR-amplified (73 °C for 26 cycles) using selective PCR primers. Libraries were then sequenced on six 100-bp single-end Illumina lanes. Raw sequencing data for the reduced-representation sequencing (nextRAD) is available through NCBI: PRJNA1248026.

Using TrimGalore v.0.6.10-1 (https://github.com/FelixKrueger/TrimGalore), Nextera adapters (at least 5 bp overlap) were removed from sequenced reads, low-quality sequences were trimmed (PHRED-quality score <20) and filtered (<30 bp reads were removed), and poor-performing samples (<40 Mb) were removed. To determine an appropriate genome for alignment, sequences were indexed and mapped against both the *A. tenuis* (Thomas et al., 2021) and *A. hyacinthus* (López-Nandam et al., 2023) genomes using BWA (Li & Durbin, 2009). Alignment to *A. hyacinthus* resulted in a higher percentage of mapped reads, thus, we mapped the sequences against this genome. iPYRAD v.0.9.94 (Eaton & Overcast, 2017) was used for mapping reads against the *A. hyacinthus* genome, locus clustering, and variant calling (85% clustering threshold, 6 as minimum coverage, minimum 4 samples per locus, and all other settings as default recommended values). The sequencing and mapping of 151 samples yielded, on average, 681,224 (± 611,783, standard deviation) mapped reads, 76,427 loci, and 562,477 SNPs.

### Population genetic structure

Sample performance was evaluated as the percentage of missing data using VCFtools (Danecek et al., 2011). Clones were identified by calculating genetic distance/similarity (Hamming-based distance) between all pairs of individuals using vcf_clone_detect.py (Bongaerts et al., 2021). The percentage of missing data and genetic distance were assessed along with the neighbor-joining tree to avoid calls from low genotyped samples (Supp. Fig. 1). Low-performance samples (>98% missing data) and variants present in <20% of the samples were removed, and a single representative individual per group was selected for groups with ≥96.5% genetic similarity. Data were filtered to only consider biallelic SNPs genotyped for >30% of individuals in each region. After filtering low-performance and outlier samples, rare variants, and potential clones (n = 11), the overall genetic structure was evaluated on 93 individuals and 124,926 loci, and a dataset of 71 individuals and 12,392 loci was used for downstream analyses on the main lineage.

Overall genetic structure was evaluated using STRUCTURE (K = 1-8, (Pritchard et al., 2000)) and SNAPCLUST (K = 1-12, (Beugin et al., 2018)). STRUCTURE was run with 10 replicates, 100,000 burn-in sweeps, and 500,000 MCMC repeats, and CLUMPP (Jakobsson & Rosenberg, 2007) was used to summarize the outcome and determine the optimal *K* value (estimators of the number of clusters, MedMeaK, MaxMeaK, MedMedK, and MaxMedK (Puechmaille, 2016)).

After identifying the main cluster, four outlier individuals were removed, and genetic structure within the main cluster was assessed using principal component analysis (function *dudi*.*pca* in adegenet package), *de novo* discriminant analysis of principal component (DAPC, adegenet package (Jombart, 2008; Jombart & Ahmed, 2011), SNAPCLUST (K = 1-8, (Beugin et al., 2018), and STRUCTURE (K = 1-8, (Pritchard et al., 2000), settings as described before). Genetic structure was further evaluated using STRUCTURE (K = 1-8, Pritchard et al., 2000) and a neutral dataset after removing outliers SNPs identified with PCadapt (K = 20, Luu et al., 2017; Privé et al., 2020). Isolation-by-distance was assessed using the Mantel test (function *mantel*.*randtest* in the package ade4 (Dray & Dufour, 2007)) to evaluate the correlation between individual genetic distances, the proportion of shared alleles (function *propShared* in adegenet, (Jombart, 2008; Jombart & Ahmed, 2011)) and geographic distances, calculated from geographic coordinates (using the *distHaversine* function, geosphere, Hijmans et al., 2024). IBD within the Western Coral Sea region was not assessed because it represents only two locations.

### Coral cover over time

Coral cover was assessed in Hernández-Agreda et al. (2022) and was reanalyzed here at the family level. Permanent quadrats (3 m x 3 m) were established at both shallow (10 m) and upper mesophotic (40 m) depths in 2012 and resurveyed following the methodology described in Bak & Nieuwland (1995). Quadrats were monitored in 2013, 2015, and 2017 for the WCS locations (Osprey BL, Osprey DT, and Osprey NW) and in 2014 and 2016 for the GBR locations (Great Detached and Yonge Reef). During the timeframe assessed in this study, two large-scale bleaching events (2016 and 2017) and three tropical cyclones (TCs) affected the study area: TC Zane (29 April - 2 May 2013, category 1), TC Ita (5 - 13 April 2014, category 5), and TC Nathan (9 - 25 March 2015, category 4). Quadrats were photographed using a Canon EOS 5DmIII with a 15 mm fisheye lens and a 1.4x teleconverter in an Aquatica underwater housing using two Sea & Sea YS-D1 underwater strobes. The benthic cover was annotated following de Bakker et al. (2017) in the CoralNet software (https://coralnet.ucsd.edu, (Beijbom et al., 2012)); identifying benthos in a 225-point grid projected onto the overview photoquadrat. Individual benthic points were followed over time, using 2012 as the reference time point. The family Acroporidae was further categorized into ‘table/corymbose/digitate/branching’ morphologies, which are characteristic of the genus *Acropora*, and ‘plating/encrusting/other’ morphologies, which represent other genera.

## Results and Discussion

The assessment of benthic coral cover revealed mostly high abundance of Acroporidae at upper mesophotic depths across five locations (Figure 1A). Nevertheless, both shallow and upper mesophotic reefs exhibited a declining trend in proportional Acroporidae cover over time, except at Great Detached, where cover of the table, corymbose, digitate, and branching morphologies remained stable (8.4 ± 0.3 %, average ± standard error). This decline in Acroporidae cover can be attributed to repeated disturbances, including two mass-bleaching events and three tropical cyclones (Harrison et al., 2019; Hughes et al., 2018; Madin et al., 2018; Marshall & Baird, 2000). Further, previous analyses of these data showed that mesophotic habitats maintained a stable coral cover through a dynamic balance between growth and mortality, with *Acropora* playing a key role as one of the fastest-growing genera, while shallow habitats exhibited declining coral cover (Hernández-Agreda et al., 2022). The dominance of Acroporidae at mesophotic depths and its overall stability over time, despite exposure to repeated disturbances, suggests that this family is likely to play an essential role as an early contributor to recovery after disturbances, as observed in shallow habitats (Linares et al., 2011; Morais et al., 2021).

**Fig. 1.**
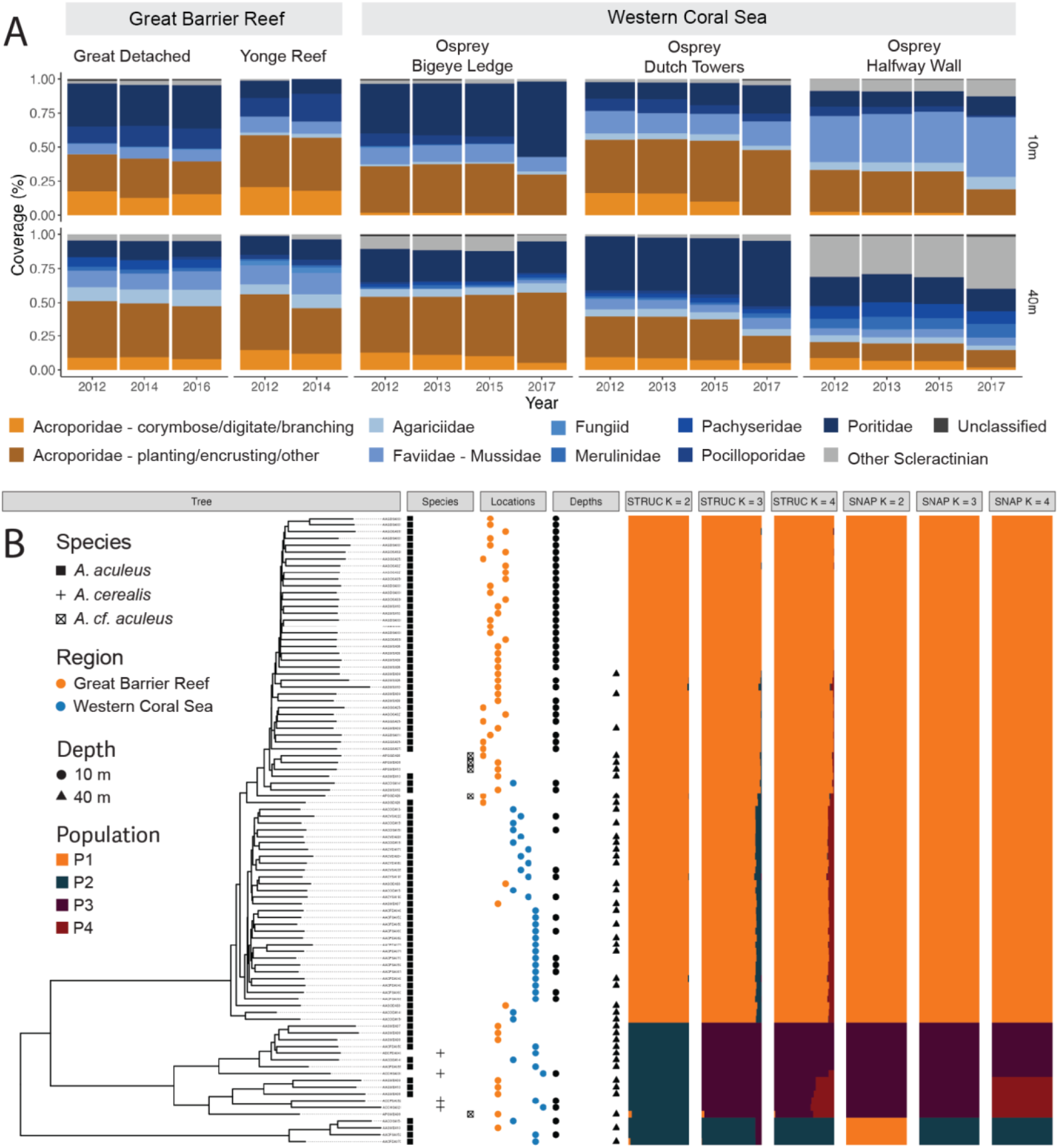
Overall genetic structure of *A. aculeus*. (A) Changes in the coral community structure at the family level in the Great Barrier Reef and the Western Coral Sea. (B) Phylogenetic tree (neighbor-joining) and genetic clustering (STRUCTURE and SNAPCLUST, *K* = 2 - 4) of collected *Acropora* specimens (based on 124,926 SNPs), with corresponding species, location, and depth.

Genetic structuring based on principal component and neighbor-joining tree analyses revealed three genetically divergent clades across the two sampled bioregions (Figure 1B). Most samples clustered together (orange in STRUCTURE *K* = 3, hereafter referred to as the main lineage) and were all identified as *A. aculeus*, including four additional *A. cf. aculeus* specimens with atypical morphology. A second smaller cluster contained the *A. cerealis* specimens (in purple in STRUCTURE *K* = 3) but also 10 specimens identified as *A. aculeus* (with one being atypical), with a further subdivision into smaller clades. The third cluster (in green in STRUCTURE K = 3) consisted of four specimens identified as *A. aculeus* but was clearly divergent from the main *A. aculeus* cluster, to which most specimens belonged. The findings highlight the ongoing challenges in *Acropora* systematics (Cowman et al., 2020; Kitahara et al., 2016; Wallace et al., 2012), and the need for an in-depth systematic assessment of these related *Acropora* species.

Clustering and ordination analyses of the *A. aculeus* main lineage revealed regional differentiation between WCS and GBR (Figure 2). Overall isolation-by-distance (IBD) was significant, but genetic dissimilarity between regions was higher than within regions (“between regions” vs. “within GBR” and “within WCS,” Supp. Fig. 2) despite these representing similar geographic distances. Only three potential first-generation migrants from the WCS were identified in the GBR, and one migrant from the GBR was observed in the WCS. Admixture signals were detected in the GBR, with several admixed individuals across all locations, whereas minimal admixture was observed in the WCS. These results suggest that gene flow primarily occurs from the WCS to the GBR, with limited flow in the opposite direction, indicating that the WCS may function as an isolated system despite occasional long-distance dispersal of *A. aculeus* larvae. Regional differentiation between WCS and GBR is consistent with the westward flow of the southern equatorial current (Steinberg, 2023) and has been previously described in other *Acropora* species (*A. millepora* and *A. kenti*, formerly known as *A. tenuis* (Bridge et al., 2024)). Restricted gene flow between these regions is likely due to the absence of reefs across the large oceanic expanse separating WCS and GBR (Lukoschek et al., 2016a).

**Fig. 2.**
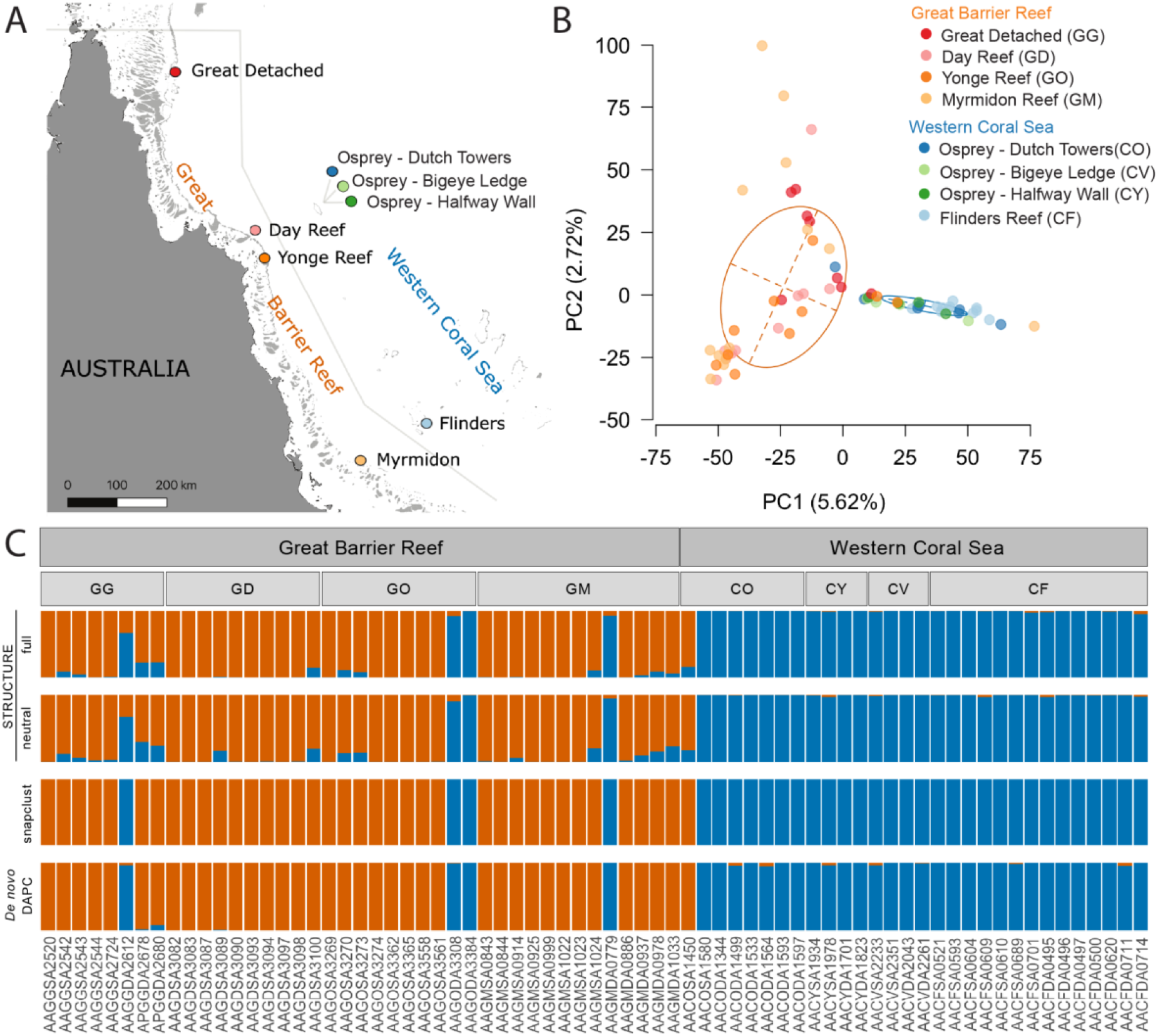
Genetic structure in the main *A. aculeus* lineage. (A) Map of sampling locations on the Great Barrier Reef and the adjacent Western Coral Sea. (B) Principal component analysis (PCA) (C) Genetic structure based on STRUCTURE results (top), snapclust (middle), and *de novo* DAPC (bottom) within the main *Acropora aculeus* lineage across regions, locations, and depths. PCA, full STRUCTURE, snapclust, and *de novo* DAPC based on 12,392 SNPs. Neutral STRUCTURE based on 11,034 SNPs.

Clustering analyses did not identify differentiation between reef locations in either region (Supp. Fig. 3), showing large dispersion and overlap among GBR locations (from far northern to central GBR) along the second axis of the PCA (Figure 2B) and non-significant IBD (Supp. Fig. 2). This pattern aligns with the genetic distribution and connectivity of *A. millepora* and *A. kenti* along the GBR. The far northern, northern, and occasionally central GBR reefs form a single cluster, while the southern reefs form a second cluster, which tends to accumulate alleles transported via the southward flowing East Australia Current (Lukoschek et al., 2016b; Matias et al., 2023; Riginos et al., 2019; Thomas et al., 2014; van Oppen et al., 2015, 2011). The boundary between these two clusters varies by species but occurs between 18°S and 21.5°S (Lukoschek et al., 2016b; Thomas et al., 2014). Although our study did not include southern reefs, the scattered distribution of Myrmidon reef samples in the PCA (Figure 2B) could indicate the presence of a latitudinal break in the *A. aculeus* connectivity near this latitude (18°S). Alternatively, the variation observed at Myrmidon may also be explained by population bottlenecks and founder effects with stochastic recruitment, as this population has experienced at least two massive mortality events in the past (11 - 75% in 1982 and 70 - 80% in 2002, (Fisk & Done, 1985; Van Oppen et al., 2011)). Despite the clear separation between the GBR and WCS regions, no further geographic genetic structuring was observed within regions.

*A. aculeus* is a depth-generalist with a broad distribution, spanning from the intertidal zone to depths exceeding 40 m (Turak & DeVantier, 2019). No genetic differentiation was detected between shallow (10 m) and mesophotic (40 m) depths when the data was analyzed collectively (Supp. Figure 3). This indicates that despite the wide depth distribution, there are no depth-associated lineages as observed for many “depth-generalist species” in the Atlantic and Pacific (Bongaerts et al., 2017; Prata et al., 2022; Rippe et al., 2021; Sturm et al., 2020; Van Oppen et al., 2018). Although this may indicate a certain potential in support of the deep reef refuge hypothesis (DRRH), the limited sample sizes (largely due to the poor sequencing results) per depth across locations constrained our ability to assess local genetic differentiation based on allele frequencies and therefore the potential and role for vertical connectivity on coral reefs.

## Conclusions

Population genomic analysis using genome-wide sequencing data has been crucial in revealing cryptic speciation and disentangling latitudinal and longitudinal connectivity patterns, which have important implications for the conservation and management of coral reefs. In this study, we uncovered the potential for cryptic diversity within *A. aculeus* but also found shallow and mesophotic populations representing the same species (with no indications of genetic differentiation over depth). Geographic assessment of the main *A. aculeus* cluster corroborated the strong genetic differentiation between the GBR and WCS regions, although there is some evidence of migrants and admixture from the WCS to the GBR. We hypothesize that mesophotic *A. aculeus* populations may be a source of larvae capable of recruiting across depths, which could be critical following acute shallow-restricted disturbances in isolated reef systems like the Western Coral Sea. With mesophotic reef habitats forming a crucial part of the reef ecosystem continuum, our results underscore the importance of further exploring biodiversity at mesophotic depths and their potential contribution to the persistence of coral reefs.

## Acknowledgments

We are grateful for the field support of David Whillas, Kyra Hay, Jaap Barendrecht, Linda Tonk, David Aguirre, Pedro Frade, Manuel Gonzalez-Rivero, David Harris, Susie Green, Erin McFadden, Steve Lindfield, and Sara Naylor, as well as Underwater Earth, The Ocean Agency, and the crews from Reef Connections, Mike Ball Dive Expeditions, SY Ethereal, and the Waitt Foundation. This project was supported, in chronological order, by the XL Catlin Seaview Survey (2012−2014; funded by the XL Catlin Group in partnership with Underwater Earth and The University of Queensland), an Australian Research Council Discovery Early Career Researcher Award (2016−2018; DE160101433), and the Hope for Reefs Initiative at the California Academy of Sciences. We also acknowledge the Waitt Foundation and the Joy Foundation for generously providing substantial sea time.

## Supplementary material

**Supp. Figure 1.**
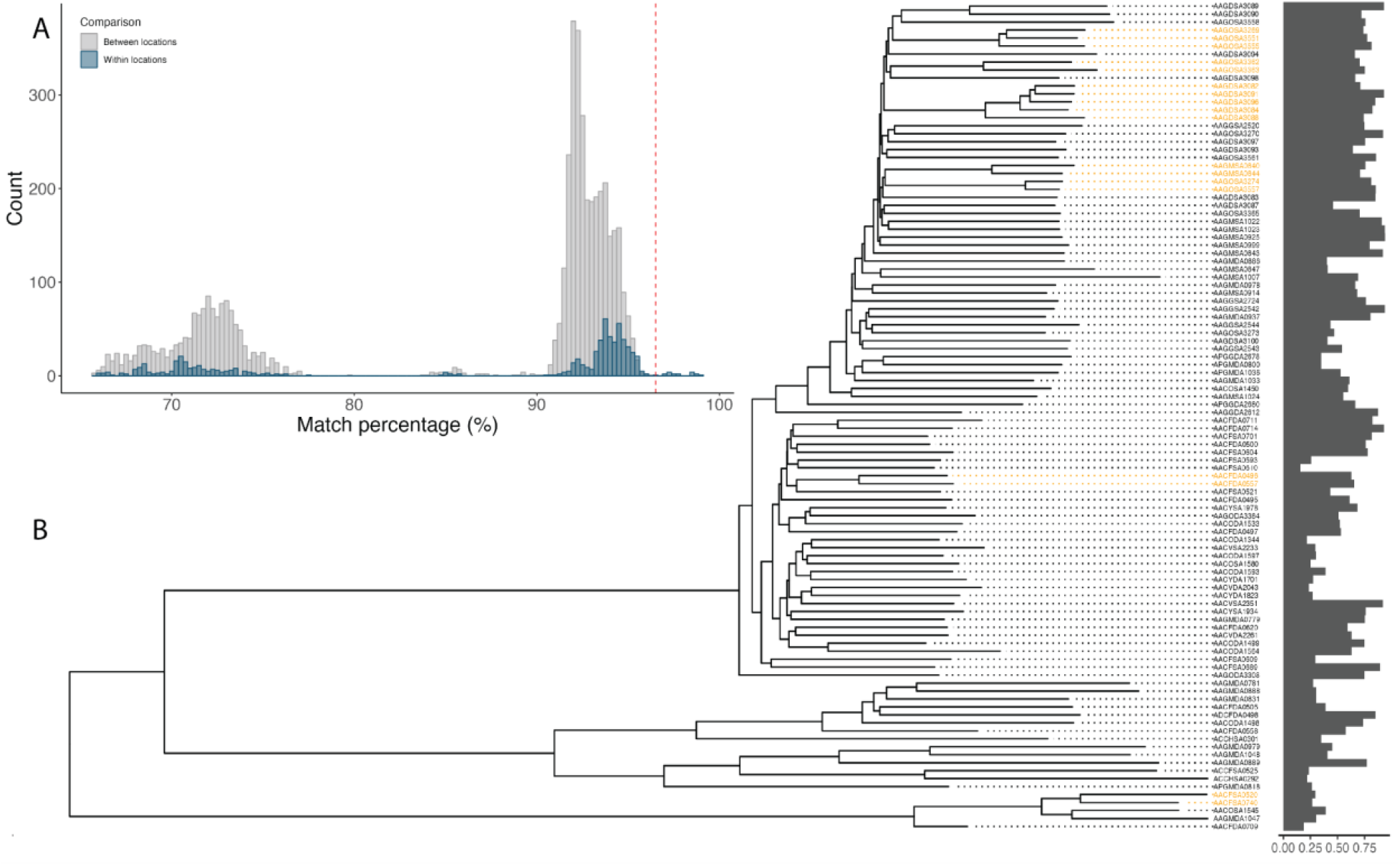
Genetic similarity and clonality threshold. (A) Histograms of pairwise genetic distance. The dashed line indicates the clonality threshold (96.5%). (B) Neighbor-joining tree and percentage of genotyping across individuals. Identified clones (similarity >96.5%) are colored in orange.

**Supp. Figure 2.**
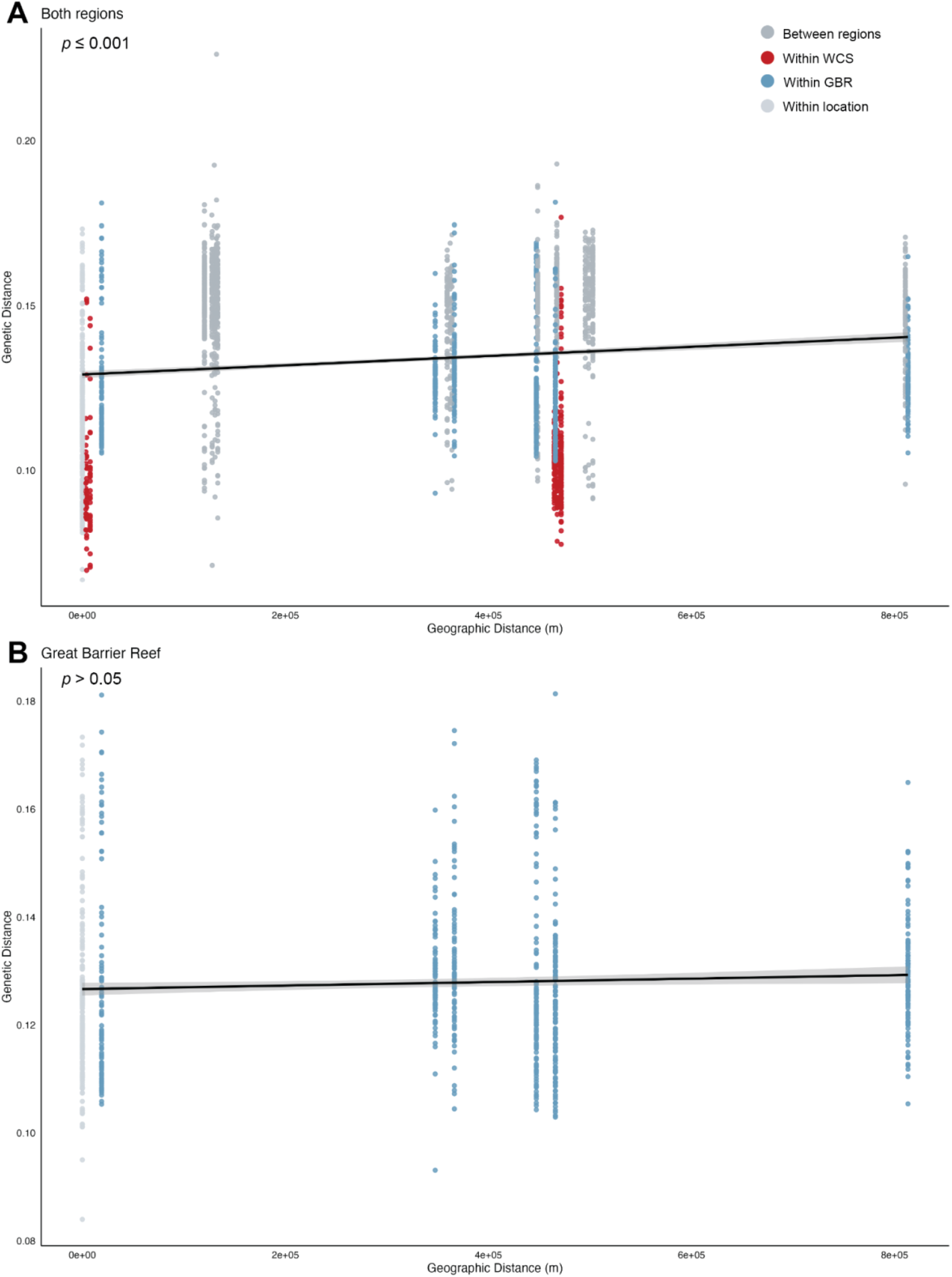
Isolation-by-distance (IBD) plots for both regions (A) and the Great Barrier Reef (B). Linear regression (in red; standard error in gray) between the genetic and geographic distances (in meters).

**Supp. Figure 3.**
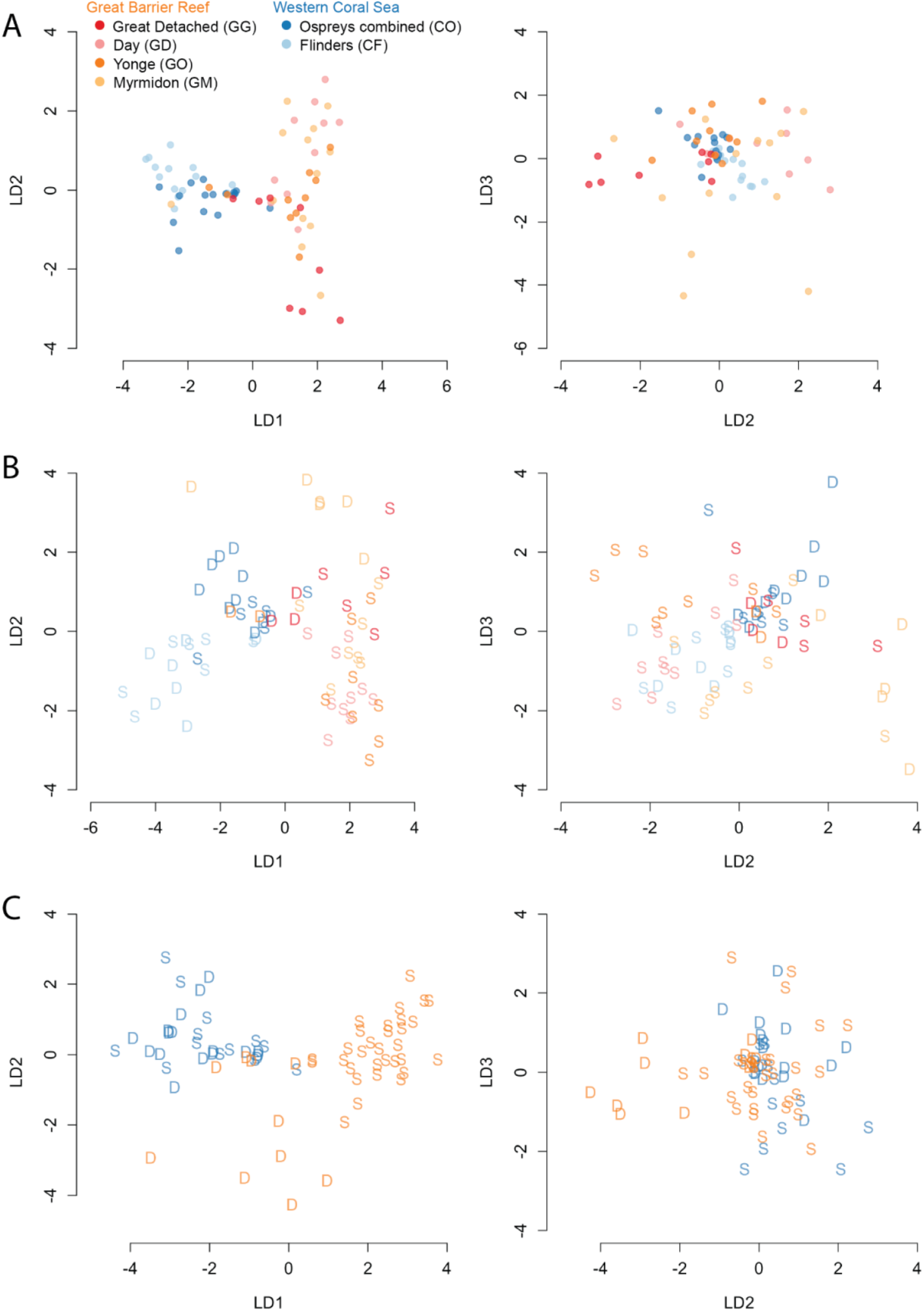
Assessment of *A. aculeus* main lineage genetic structure. Discriminant Analysis of Principal Components (DAPC) for the interactions (A) regions and locations, (B) regions, locations, and depths, and (C) regions and depths. Colors of sampling depths, locations, and regions are indicated in the inset in (A).

**Table S1.** Estimators of the number of STRUCTURE clusters. Main lineage data refers to Figure 2.

| K | Main lineage (full) STRUCTURE analysis |  |  |  | Main lineage (neutral) STRUCTURE analysis |  |  |  |
| --- | --- | --- | --- | --- | --- | --- | --- | --- |
|  | Med<br>MeaK | Max<br>MeaK | Med<br>MedK | Max<br>MedK | Med<br>MeaK | Max<br>MeaK | Med<br>MedK | Max<br>MedK |
| 2 | 2.0 | 2 | 2.0 | 2 | 2.0 | 2 | 2.0 | 2 |
| 3 | 2.0 | 3 | 2.0 | 3 | 2.0 | 3 | 2.0 | 3 |
| 4 | 3.0 | 3 | 3.0 | 3 | 2.0 | 4 | 2.0 | 4 |
| 5 | 2.0 | 3 | 2.0 | 3 | 2.0 | 2 | 2.0 | 2 |
| 6 | 2.0 | 3 | 2.0 | 3 | 2.0 | 4 | 2.0 | 4 |
| 7 | 2.0 | 3 | 2.0 | 3 | 2.0 | 2 | 2.0 | 2 |
| 8 | 2.0 | 3 | 2.0 | 3 | 2.0 | 2 | 2.0 | 2 |

## Notes

https://www.ncbi.nlm.nih.gov/bioproject/PRJNA1248026/

### Competing Interest Statement

The authors have declared no competing interest.

